# Uncoupling of Chromatin Assembly from DNA Replication in *Sciara* Reveals a Domain of Postreplicative Immature Chromatin

**DOI:** 10.1101/2021.05.12.443896

**Authors:** Fyodor D. Urnov, Ulrich Scheer, Hanswalter Zentgraf, Heidi S. Smith, Susan A. Gerbi

## Abstract

DNA replication in dividing eukaryotic cells imposes a requirement for the faithful recreation on the newly synthesized chromatids of the nucleoprotein architecture of parent chromosomes. Practically nothing is known about the structure of postreplicative immature chromatin—a very short-lived entity (< 30 min.). We report here the unexpected discovery that during DNA amplification of locus II/9A in salivary gland polytene chromosomes of the fungus fly *Sciara coprophila*, DNA replication fork passage is uncoupled from postreplicative chromatin assembly; this enables visualization and analysis of chromatin fibers disassembled by DNA replication. We used electron microscopy to visualize a wealth of low nucleosome density immature chromatin fibers in preparations of *Sciara* chromatin from amplification-stage tissue. Remarkably, as gauged by high sensitivity to micrococcal nuclease and an unusually short length of DNA associated with each histone octamer, we found that locus II/9A which undergoes amplification and is replicated once every 4-6 hrs.—but not the bulk genome or a replicatively quiescent DNA stretch—was maintained in such an ummature fiber for ca. 24 hrs. Following amplification, locus II/9A assumed conventional chromatin organization, indicating that the epigenetic mark targeting nascent DNA to the chromatin assembly machinery is stable for several hours. We propose that this very unusual prolonged maintenance of a segment of the genome in immature chromatin facilitates access by the basal transcriptional machinery to the amplified DNA, and thus is an evolutionary adaptation to the demand for high transcription from genes that reside in the amplified loci.

## INTRODUCTION

In proliferating eukaryotic cells, the continuity of genomic function relies not only on the accuracy of DNA replication during S phase, but also on the faithful recreation of the nucleoprotein architecture of the parental chromosomes on daughter chromatids (1). In contrast to DNA replication, however, the details of how chromatin replicates remain elusive. Is it known that a moving DNA replication fork disrupts both histone-DNA contacts (2, 3) as well as nonhistone protein-DNA complexes (4), and that nascent DNA rapidly (on the scale of minutes) becomes associated with proteins in a poorly understood interplay between three processes: random segregation of parental histones onto daughter chromatids (5), pre-emptive binding of nonhistone regulatory factors to their target sites in the genome (6), and sequential (first H3/H4, then H2A/H2B) *de novo* deposition of hyperacetylated histones onto DNA by dedicated chaperones (7).

Once deposition is complete, chromatin is rapidly (< 30 min.) converted into a mature state by histone deacetylases (8). In addition to deacetylation, chromatin maturation involves an active reorganization of histone-DNA contacts to ensure that the newborn nucleosomes are evenly spaced. Extensive biochemical analysis from many laboratories revealed that the eukaryotic nucleus is populated with a variety of ATPase-containing complexes (9)—e.g., NURF (10), ACF (11), Mi-2 (NuRD/NRD) (12, 13), and CHRAC (14)—all of which can act *in vitro* to effect histone octamer mobilization and thereby convert disorganized nucleosomal arrays into a fiber with regular octamer spacing.

Considering the wealth of interest in this subject, it is remarkable that the chaperones, the HDACs, and the ATPases responsible for chromatin assembly and maturation *in vivo* all remain unidentified. In fact, our understanding of the structure of postreplicative immature chromatin remains confined to its operational definition first proposed 25 years ago, when it was discovered that newly synthesized DNA in proliferating cells is more sensitive to nucleases than bulk genomic DNA (15). To some extent this paucity of information is inevitable, since immature chromatin is a very short-lived entity: histones are deposited on the DNA within minutes of replication fork passage and their deacetylation takes only 20-30 minutes (7). In addition, replication fork progression in nuclei of metazoan cells is asynchronous, and the nature of the replication origins where they initiate remains poorly understood (16, 17). Thus, it has not yet been possible to obtain a population of cells in which a given locus would be in the 30 min. postreplicative window at the time of the experiment.

We describe here our unexpected discovery of a prolonged uncoupling of replication fork passage from chromatin assembly in the lower dipteran, *Sciara coprophila*, and exploit this finding to directly visualize, localize to a specific stretch of the genome, and analyze the heretofore elusive postreplicative immature chromatin.

An interesting feature of *Sciara* biology offers a unique opportunity to investigate postreplicative chromatin maturation: locus- and tissue-specific DNA amplification. Late in larval development, *Sciara* salivary glands produce significant quantities of protein for the pupal cocoon. High demand for specific transcripts is met by intrachromosomal amplification of the cognate DNA segments at a precise time in development, while the bulk of the genome is completing a single round of replication. Thus, specific DNA replication origins (ORIs) fire repeatedly to generate excess template for transcription. For example, locus II/9A contains genes that code for components of the cocoon (18), undergoes an additional 3-4 rounds of replication (19), and is overrepresented ca. 17-fold in the genome (20). Once amplification is complete, all DNA synthesis in the salivary gland ceases; the ensuing burst of transcription from the amplified loci is accompanied by massive localized swelling to form the “DNA puffs” on the polytene chromosomes, and later is followed by salivary gland apoptosis.

In the present study, we used electron microscopy to reveal a variety of replicative intermediates and immature nucleosomal fibers in amplification-stage *Sciara* salivary gland chromatin preparations. Remarkably, molecular probes revealed the very unusual persistent locus-specific maintenance of immature chromatin over a stretch of the genome undergoing DNA amplification, and a ca. 24 hr. lag period before a mature nucleosomal array was assembled. Since a round of replication in this locus occurs once every 6-8 hrs., such extraordinary delay of chromatin maturation kinetics represents the first characterized instance of an *in vivo* uncoupling of postreplicative chromatin assembly and maturation from replication fork passage. We hypothesize that the persistent maintenance of the amplified loci in immature chromatin faciliates access by the basal transcriptional machinery, and thus may be an evolutionary response to the demand for high mRNA synthesis by genes found in the amplified DNA stretches.

## MATERIALS AND METHODS

### Chromatin spread preparations

*Sciara coprophila* (Insecta: Diptera: Nematocera) salivary glands were dissected from larvae at the peak of amplification activity (“12×6” as per the eyespot index (20, 21)) and from post-amplification stage larvae (“edge-eye”) and incubated in low salt buffer (0.1 mM Na borate, pH 8.5-9.0) containing 100 μg/ml tRNA and 0.03% Sarkosyl. Nuclei were manually collected from the ruptured cells and transferred into a droplet of the same buffer placed on a siliconized glass slide at 4° C. After 20 min. the dispersed nuclear contents were centrifuged through 1% formaldehyde (prepared from paraformaldehyde), 0.1 M sucrose, 0.1 mM Na borate buffer (pH 8.5) onto a freshly glow-discharged, carbon-coated grid. After centrifugation, the electron microscope grid was briefly immersed in 0.4% Kodak Photo-flo, air-dried, stained in ethanolic 1% phosphotungstic acid, dehydrated in 100% ethanol and air-dried again. Finally the preparation was rotary-shadowed with platinum-palladium (80:20) at an angle of ca. 7° (for a detailed protocol of the Miller chromatin spreading technique, see ref. (22)). Micrographs were taken with a Zeiss EM10 electron microscope.

### Micrococcal Nuclease Probing of Chromatin

Twelve amplification stage or 8 post-amplification stage pairs of salivary gland pairs were manually dissected in CR medium (87 mM NaCl, 3.2 mM KCl, 1.3 mM CaCl_2_, 1 mM MgCl_2_, 10 mM Tris-Cl pH 7.3 (23)) and then transferred into 100 μl of ice-cold buffer “C” (15 mM Tris pH 7.5, 60 mM KCl, 15 mM NaCl, 10 mM NaHSO_3_, 0.34 M sucrose, 0.15 mM β-mercaptoethanol (24)). Nonidet-P40 was added to a final concentration of 0.5% (v/v), and the salivary glands were ruptured by gently homogenizing on ice, followed by a 15 min. incubation ice with gentle mixing every 3 minutes. Nuclei were released from salivary gland cells by passing the sample through a tapered P-20 pipetman tip. The samples were warmed to 37°C for 1 minute, and then treated with 6 Kunitz units/ml of MNase (NFCP grade, Worthington) for 2 minutes at 37°C. The reaction was stopped by adding an equal volume of buffer TNESK (20 mM Tris pH 7.4, 200 mM NaCl, 2 mM EDTA, 2% SDS, and 200 μg/ml proteinase K (25)); the samples were then incubated at 37°C overnight. DNA was extracted once with an equal volume of buffered phenol (Amresco), once with a mixture of phenol/chloroform/isoamyl alcohol (50:48:2--v:v:v), and once with chloroform. RNA was then removed by treating with 5 μg/ml RNase A for 15 minutes, followed by another round of phenol-chloroform extraction, and ethanol precipitation. Following precipitation, DNA was resuspended in 1x TE buffer (10 mM Tris-Cl, pH 8.0, 1 mM EDTA);

Five DNA samples prepared this way were pooled to prevent sample-to-sample variation; one-fifth of each sample was then electrophoresed on a 2% high-resolution blend agarose (Amresco) gel in 1x Tris-acetate-EDTA buffer (40 mM Tris-acetate, 1mM EDTA). The gel was processed, transferred to maximum strength Nytran Plus membranes (Schleicher and Schuell), hybridized with DNA probes labelled by the random-priming procedure (Boehringer Mannheim), and washed according to the manufacturers’ instructions, except that no depurination was performed to prevent loss of mononucleosome-size DNA fragments; all washes were performed at 65°C, and the stringency of the final wash was 0.1x SSC and 0.5% SDS. Filters were air-dried briefly, wrapped in Saran Wrap, and exposed to Konica general purpose X-ray film for 0.5-10 days at -70°C with intensifying screens or to Bio-Imaging Analyzer plates.

## RESULTS

Chromatin from *Sciara* salivary gland nuclei was visualized by EM; to confirm that the preparation methods preserve the nucleosomal organization of chromatin, we analyzed samples from replicatively quiescent post-amplification tissue. In agreement with expectation, the overwhelming majority of structures visualized (Fig. 1A-B) were conventional “beads-on-a-string” nucleosomal fibers (26).

**Fig. 1.**
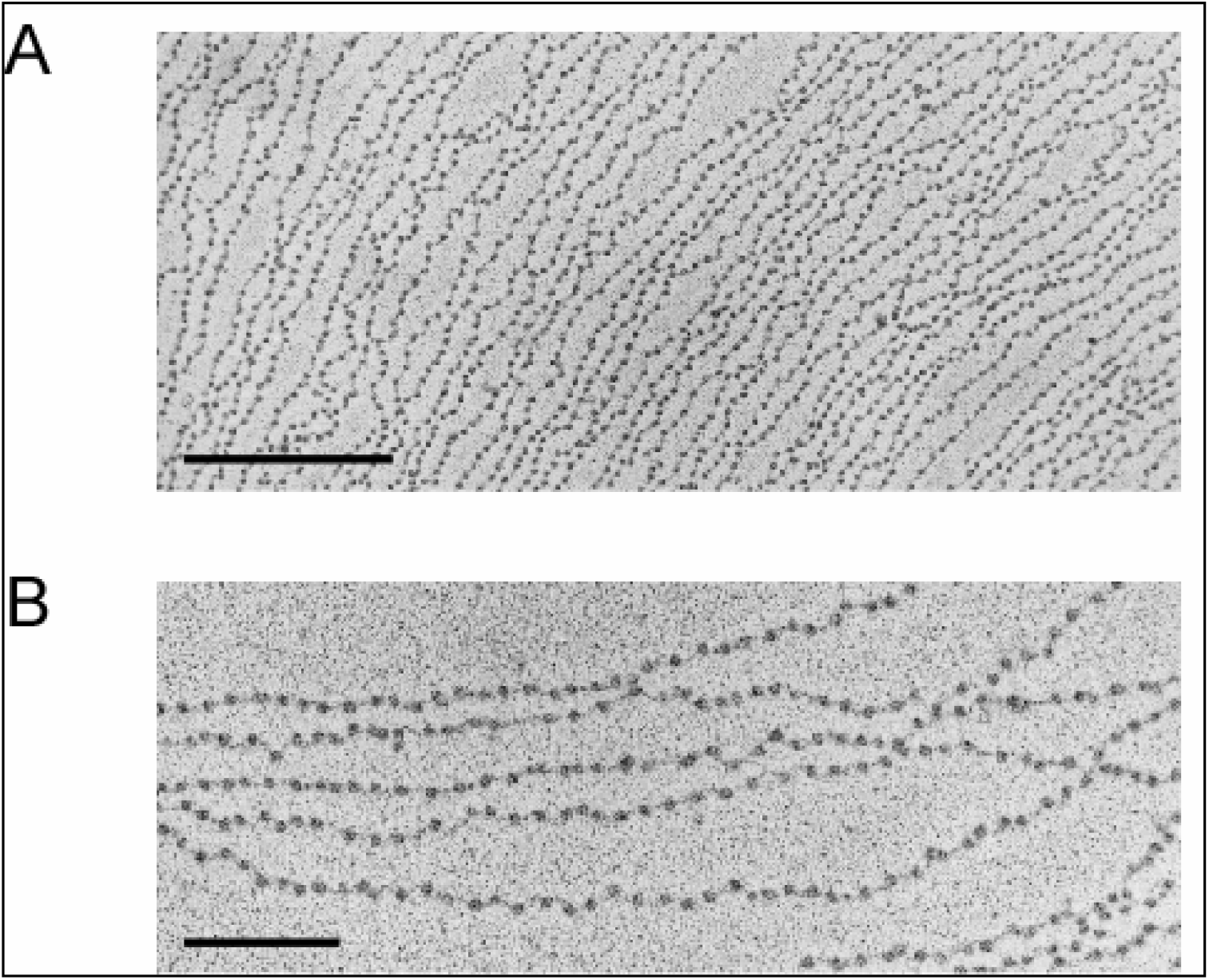
Conventional nucleosomal organization of *Sciara* chromatin during post-amplification stages. Preparations of post-amplification stage *Sciara* chromatin were analyzed by EM *(A)*; magnification is 35,000x (bar = 0.5 μM). A portion of panel A is shown magnified in panel B (magnification is 63,000; bar = 0.2 μM).

Preparations from salivary glands in which the DNA puff loci were undergoing amplification (i.e., repeated rounds of replication), contained fibers of intact chromatin (Fig. 2B-1 and 2C-6), and a a variety of replicative intermediates: (i) replication bubbles of various size in which both arms appeared coated with electron-dense material (Fig. 2B-2/3); (ii) replication forks in which one strand appeared significantly less electron-dense than the other (Fig. 2B-4). The nature of the electron-dense material is unknown, but we note that these structures are strikingly similar to replicative intermediates seen on viral genomes in herpes simplex (27, 28) and adenovirus-infected cells (29).

**Fig. 2.**
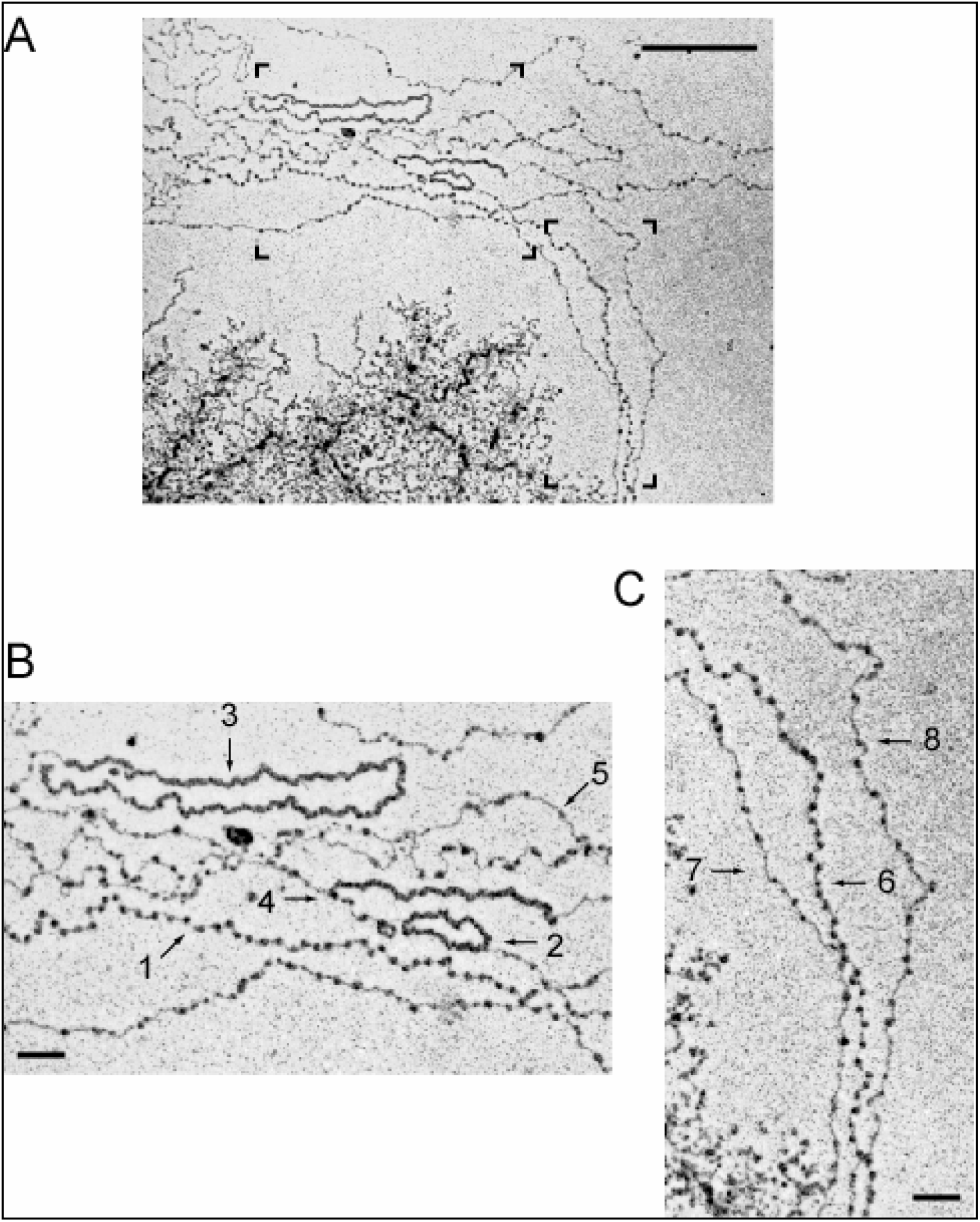
Multiple replication intermediates and immature chromatin fibers in chromatin samples from amplification stage *Sciara* salivary glands. Preparations of amplification stage *Sciara* chromatin were analyzed by EM *(A)*; magnification is 30,000x (bar = 0.5 μM). The left bracket in panel A is magnified in panel B, and the right—in panel C (magnification in both panels is 60,000x; bar = 0.1 μM). Numbered arrows indicate characteristic chromatin structures seen in this analysis (see text). The rDNA “Christmas trees” can be seen in the lower left-hand corner.

Considering that conventional chromatin assembly and maturation is known to occur during a short window of time (7), we were surprised to find a significant number of incompletely assembled (immature) chromatin fibers in preparations from amplification-stage tissue (Fig. 2A-C). These contained extensive stretches of seemingly protein-free DNA interspersed with bead-like particles, many of which appeared smaller and less electron-dense that nucleosomes visualized on immediately adjacent mature fibers (compare Fig. 2C-7/8 with 2C-6). We also observed fibers that appeared largely protein-free (Fig. 2B-5).

DNA amplification of specific loci in the *Sciara* genome occurs when the rest of the genome is completing its final round of replication. To determine if the unusual nucleoprotein fibers seen in the electron micrographs were derived, in part, from amplifying loci, we used micrococcal nuclease (MNase) to probe chromatin assembled at segment 9A on *Sciara* chromosome II (the location of one of the earliest and largest DNA puffs). A 14 kb fragment of the II/9A locus contains a 1 kb major DNA amplification ORI (30) located 2.5 kb upstream of gene II/9-1 whose product is thought to form part of the pupal cocoon (18). To distinguish passive disruption of histone-DNA contacts that occurs due to DNA replication from active localized chromatin remodeling targeted by regulators of replication and transcription to this area, we made use of a DNase I hypersensitive site (DHS) located next to the 1 kb replication ORI (Fig. 3A and data not shown).

**Fig. 3.**
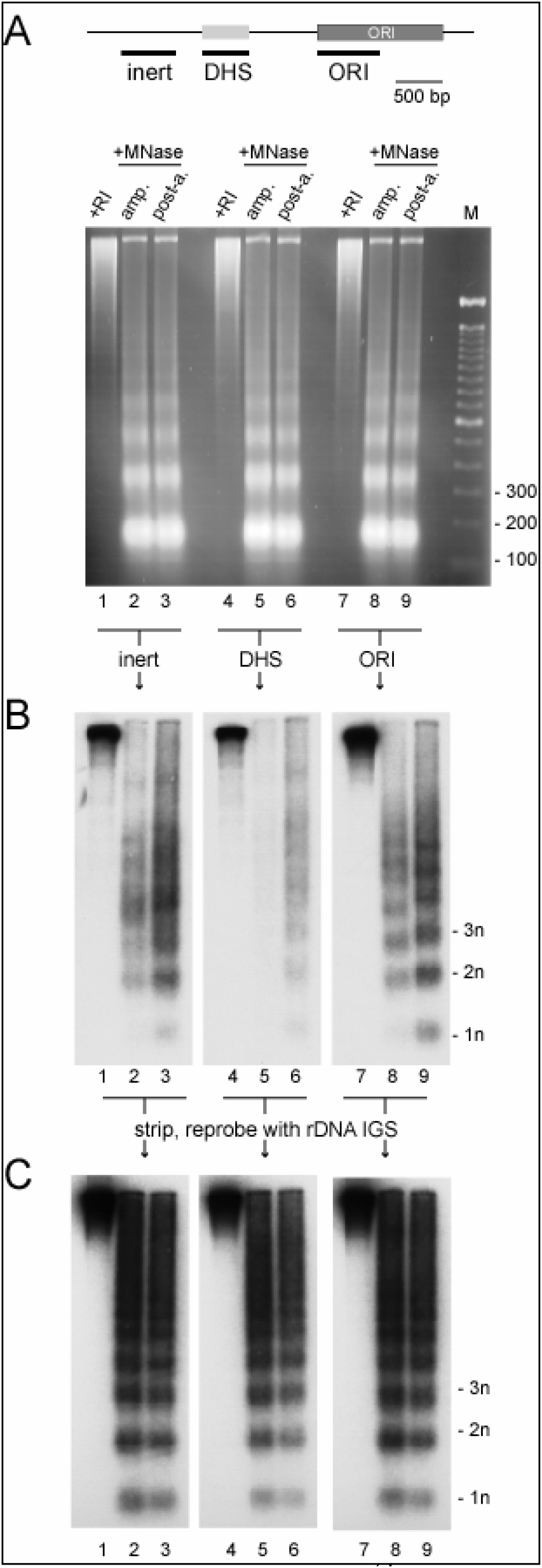
Low nucleosomal density and shortened length of DNA in nucleosome core particle in II/9A DNA during amplification. Amplification (lanes 2, 5, 8) and post-amplification (lanes 3, 6, 9) chromatin samples were treated with MNase and resolved on a high-percentage agarose gel alongside an aliquot of *Sciara* genomic DNA digested with EcoRI (lanes 1, 4, 7) *(A)*. Following transfer to a nylon membrane, lanes 1-3 were probed with a 400 bp fragment located upstream of the DHS (“inert”), lanes 4-6—with the DHS, and lanes 7-9—with the ORI *(B)*. The positions of mono-, di, and trinucleosomal particles are indicated. The membranes were then stripped and all three were reprobed with a fragment of the rDNA IGS *(C)* to prove that the three membranes contained identical DNA preparations.

MNase exhibits marked preference for DNA in the linker between adjacent nucleosomes, and thus is commonly used to measure nucleosome density—the number of histone octamers associated with a given DNA stretch (31). We treated *Sciara* salivary gland nuclei with MNase; a ladder of mono-, di-, and higher-order nucleosomal particles were visualized in chromatin (Fig. 3A). As expected, DNA amplification (a localized phenomenon) did not affect global nucleosomal organization of the *Sciara* genome, and amplification and post-amplification nuclei yielded identical nucleosomal ladders (lanes 2-3, 5-6, 8-9 in Fig. 3A). To obviate concerns about DNA loss during sequential stripping and reprobing of nylon membrane, we prepared a large sample of chromatin, split the resulting DNA into multiple aliquots, and resolved them on the same gel (Fig. 3A). Thus, in Fig. 3A, DNA in lane 2, 5, and 8 come from the same sample of amplification-stage DNA, and in lanes 3, 6, and 9—from the same sample of post-amplification stage DNA. To adjust for the specific activity of the probe and its hybridization efficiency, we included an identical aliquot of EcoRI-digested *Sciara* genomic DNA on each membrane (lanes 1, 4, and 7). We then used short DNA probes derived from the II/9A locus (Fig. 3A) in Southern blot analysis.

As expected, DNA over the DHS was very sensitive to MNase (Fig. 3B, lane 5), and was rapidly degraded by the nuclease; extensive exposure of the membrane failed to reveal a nucleosomal ladder over the DHS. Surprisingly, during DNA amplification, other fragments in the II/9A locus also exhibited a nucleosomal organization markedly different from that of the bulk genome. For example, a randomly chosen stretch of DNA outside of the DHS (“inert” probe) had low nucleosome density (Fig. 3B, lane 2), as did the ORI (Fig. 3B, lane 8) when compared to the standard nucleosomal profile of the bulk genome at amplification stage (Fig. 1, lanes 2 and 8). This difference was particularly striking when these same filters were reprobed with a replicatively quiescent fragment of the *Sciara* genome (the rDNA intergenic spacer, IGS (32); Fig. 3C) to reveal the robust assembly of amplification-stage rDNA into mature chromatin (Fig. 3C, lanes 2, 5, 8 in Fig. 3C).

In addition to low density of histone octamers over II/9A DNA, nucleosomes found in this locus assumed an altered conformation: comparison of replicating vs. quiescent chromatin showed that the MNase-resistant di- and trinucleosomal particles were noticeably smaller during DNA amplification (compare location of “2n” and “3n” fragments in Fig. 3B, lanes 8 and 9, or lanes 2 and 3). This peculiarity was specific to the amplifying DNA stretch, since the rDNA (Fig. 3C) and the bulk genome (Fig. 3A) did not reveal a difference in di- and trinucleosome size between amplification and post-amplification samples. We analyzed other fragments of the II/9A locus and also observed low nucleosomal density and unusual brevity of the nucleosomal repeat during amplification (data not shown).

Importantly, in contrast to the low density of nucleosomes found at the II/9A locus during amplification stage, a standard nucleosomal ladder was observed at this locus after DNA amplification had ceased and no DNA replication was ocurring (Fig. 3B, lanes 3, 6, 9). The II/9A nucleosomal profile in post-amplification stage samples mirrored that seen over the rDNA (Fig. 3C, lanes 3, 6, 9) and in the bulk genome (Fig. 3A, lanes 3, 6, 9). Therefore, the low density of nucleosomes at locus II/9A is found during repeated rounds of DNA replication in this region, but a conventional nucleosome profile is restored after DNA amplification ceases.

## DISCUSSION

Electron micrographs of amplification-stage *Sciara* chromatin revealed a number of postreplicative intermediates, as well as incompletely assembled chromatin fibers (Fig. 2). These novel structures may represent postreplicative immature chromatin, because neither type of structure could be seen in preparations from nonreplicating cells (Fig. 1). Its two distinguishing characteristics are a low nucleosome density, likely caused by dilution of parental histones during replication, and the appearance of “subnucleosomal particles,” which may represent a chromatin assembly intermediate, such as the histone H3/H4 tetramer complexed with ∼130 bp of DNA. Molecular analysis of DNA over an amplifying locus revealed that during DNA replication, this DNA is found in such immature chromatin: the nucleosome density is very low, and the length of the DNA associated with a histone octamer is shorter than in mature chromatin (Fig. 3B); once amplification is complete, II/9A DNA assumes normal nucleosomal organization. Control experiments indicate that the rDNA (Fig. 3C) and the bulk of the genome (Fig. 3A), are assembled into conventional chromatin at all stages.

DNA amplification of the II/9A locus takes ca. 24 hrs., and ca. 3-4 DNA replication initiation events occur during this time; the average DNA stretch, therefore, encounters a passing DNA replication fork once every 6-8 hrs. Phenotypic markers used to stage *Sciara* larvae offer no indication as to when the puff II/9A ORI has fired, only that amplification is ongoing, and each chromatin sample in Fig. 3 contains an aliquot of DNA from ca. 500 individual larvae. Simple statistical considerations, therefore, indicate that the overwhelming majority of the DNA from locus II/9A in our preparations is not in the 30 minute post-replicative window at the time of analysis. Thus, our EM (Fig. 1) and molecular (Fig. 3) analysis both indicate that during amplification, chromatin assembly and maturation in *Sciara* locus II/9A occur with significantly (ca. 10-fold) delayed kinetics relative to the 30 min. window observed in metazoan cells.

It is possible that the failure of II/9A DNA to assemble into conventional chromatin during amplification is due to histone biosynthesis lagging behind that of the DNA. In addition, chromatin assembly chaperones and/or nucleosome spacing engines may be transiently disabled (33) during DNA amplification. The *Sciara* system is well suited for biochemical and cytological analysis of these issues; it will also be of interest to determine if amplifying loci elsewhere in the *Sciara* genome are maintained in immature chromatin.

Whatever the mechanism whereby assembly is transiently inhibited, it is remarkable that following amplification, the nascent DNA is assembled into mature chromatin (Fig. 1A and 3B). Thus, histone deposition and histone octamer spacing occurs *in vivo* on quasi-naked DNA fibers (e.g., Fig. 1D-5 and 1E-7). Newly replicated DNA in human cell extracts (34) and in budding yeast (35) is epigenetically marked by PCNA in the aftermath of replication fork passage such that the chromatin assembly machinery then interprets this tag as a targeting signal. If such a tagging system occurs in *Sciara*, then our data suggest that the epigenetic mark can remain stable over many hours.

Chromatin assembly has a well established role in preventing access of the basal transcription machinery to DNA (36-38). Genes coding for saliva proteins in *Sciara* are programmed to generate extraordinary quantities of mRNA over a short timespan. We propose that, in addition to DNA amplification that provides extra template for this transcriptional burst, the persistent failure of the amplified loci to assemble mature chromatin may be an auxiliary evolutionary adaptation to the demand for high mRNA synthesis, and that it serves to facilitate the access of the basal transcriptional machinery to regulatory DNA elements in the amplified loci. It will be of interest to determine if amplifying DNA loci in other organisms, such as the chorion loci in *Drosophila* (39, 40), exploit this auxiliary mechanism.

## ACKNOWLEDGEMENTS

We thank Ken Zaret for experimental advice on MNase treatment of chromatin. This research was supported by NIH GM35929 and NIH GM121455 to S.A.G.

## SUPPLEMENTARY FIGURE LEGEND

**Supplementary figure.**
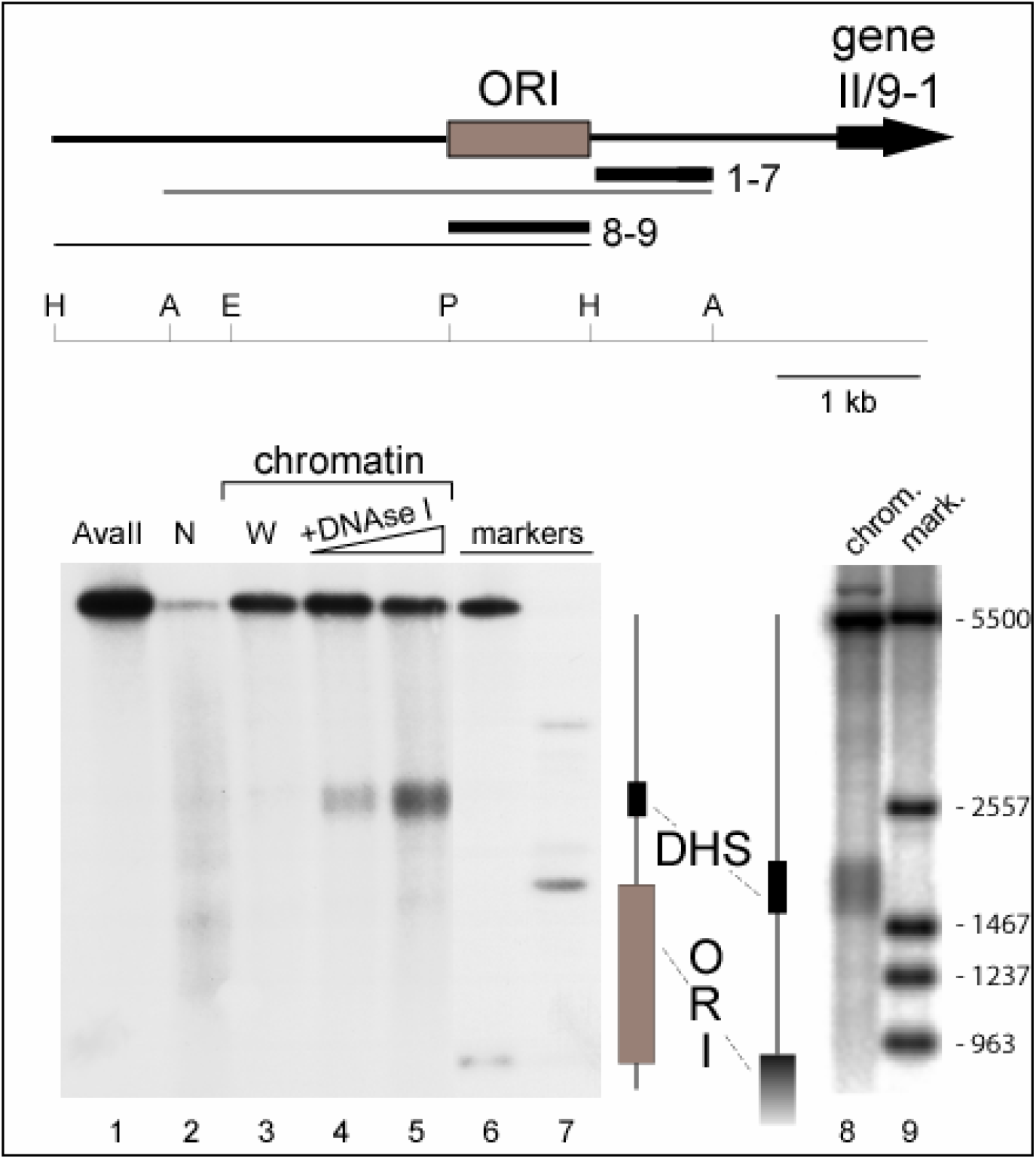
Only a small region of the II/9A locus is stably remodeled into a DNase I hypersensitive site. A map of the locus is shown; the location of the major DNA amplification origin (ORI) and the transcription unit (gene II/9-1) is indicated. The fragments analyzed by indirect end-labelling are shown as thin lines, and the probes used for Southern blotting—by thick lines; the numbers refer to the lanes in which the data are shown. A restriction map of the locus is shown below (A=AvaII; H=HindIII; P=PvuII; E=EcoRI). DNA untreated with DNAse I (lane 1), naked DNA treated with DNAse I (lane 2), nuclei that were processed in the absence of DNAse I (lane 3) are included as controls. Lanes 4-5 contain amplification stage chromatin samples treated with DNAse I, and lanes 6-7 contain genomic DNA markers digested with AvaII and an additional enzyme(s) (HindIII, and PvuII + EcoRI, respectively). Lane 8 contains a chromatin sample treated with DNAse I resolved on a high-percentage agarose gel alongside genomic DNA markers (lane 9).

